# A taxonomically informed DNA reference library to facilitate future biodiversity assessments and monitoring: a case study using seaweeds along a tropical-temperate transition zone in South Africa

**DOI:** 10.1101/2023.09.14.557690

**Authors:** Maggie M. Reddy, Jamie du Plessis, Robert J. Anderson, Rouvay Roodt-Wilding, John J. Bolton

## Abstract

The role of seaweeds in the blue bioeconomy has stimulated research efforts around the world but proper species identification and biodiversity assessments, remain a challenge. The South African coast hosts the confluence of the Indian Ocean and Atlantic Ocean, creating a dynamic evolutionary environment that has over time yielded a rich diversity of seaweeds with the highest seaweed diversity occurring along the Agulhas Marine Province. Although South Africa harbours one of the richer seaweed floras in the world, only 24% of the known species are represented by DNA barcodes. We therefore initiated the construction of a taxonomically guided DNA reference library for seaweeds in South Africa with the aim of continuously adding to it in the future. To do this, a seaweed biodiversity survey of the Rhodophyta occurring along a temperate-tropical biogeographic transition zone situated within the Agulhas Marine Province (AMP) in South Africa was carried out. Seaweeds were identified in the field using available field or taxonomic guides and herbarium vouchers were prepared. Subsamples were preserved for DNA analyses and three DNA barcodes (LSU D2-D3; *rbc*L-3P; COI 5P) were amplified. Sequences were verified on BLAST and preliminary phylogenetic analyses or comparison with the literature were carried out where necessary. A total of 220 barcodes was generated for 88 species and one species variety, including 17 species from or near their type localities and eight generitypes. Novel barcodes were generated for 73 species, nearly half of which were species endemic to Southern Africa. In addition, 21 taxa representing new, potentially new, or reinstated species and at least two new genera were identified as well as one new distribution recorded, all of which require further study. This study significantly adds to the foundational biodiversity knowledge of the South African seaweed flora and highlights new avenues for further research.

## Introduction

Algae form the base of most ecological food webs in our oceans and in turn support most commercial marine fisheries (Chapman 2013). More recently, the role of macroalgae in the blue bioeconomy has stimulated both research and seaweed awareness around the world (Ligtvoet *et al*. 2019). Although the utilization of seaweeds began centuries ago, relatively few species have been commercially exploited (Zemke-White & Ohno 1999). This is partly due to a lack of foundational biodiversity knowledge such as species inventories, biorepositories linked to commercial applications (Reddy *et al*. 2021) and reliable species identification.

Many seaweeds are difficult to identify because of their simple morphology, environmentally induced phenotypic plasticity, poorly documented life history stages, or a general lack of knowledge on reproductive morphology which may be diagnostic for some taxa (Saunders 2005). Further challenges arise when complex evolutionary processes such as convergent evolution and cryptic species are considered (Reddy 2018, Diaz-Tapia *et al*. 2020). Naturally the application of DNA has become routine in seaweed biodiversity research. DNA barcoding is a method of identification based on the premise that unique genetic identifiers can be used to catalogue and subsequently identify species against a reference library of known species (Hebert *et al*. 2003a, b). However, the utility of DNA barcoding relies on an extensive collection of well curated (Wilson *et al*. 2018) and taxonomically verified reference sequences, the availability of which tends to vary among biological organisms. For instance, model organisms or well-studied organisms tend to be better represented in terms of barcodes compared with non-modelled organisms, especially for marine organisms. The mitochondrial COI region is considered the gene of choice for DNA barcoding in most eukaryotes but is prone to artefacts as a result of hybridisation, ancestral polymorphism, incomplete lineage sorting and introgression (Moritz & Cicero 2004). To avoid these and other challenges, a multi-marker (nuclear, chloroplast, mitochondrial) approach has been adopted whereby unlinked gene regions are applied to a set of specimens, making the process more robust.

DNA barcoding surveys of seaweeds have typically adopted one of two strategies. Either a specific taxonomic group is targeted from within their distribution range (for practical reasons) or from certain geographic regions. Alternatively, all the seaweeds (or a certain group, e.g. the Rhodophyta) in a specific geographic locality or region are targeted. For example, Miladi *et al*. (2018) carried out a DNA barcoding survey of the genus *Ulva* in Tunisia, while Steinhagen *et al*. (2018) studied the families Ulvales and Fucales from Kiel Germany and Saunders & McDonald (2010) focused on the order Rhodymeniales in Australia. Other studies have targeted all three major lineages (the Rhodophyta, Chlorophyta and Phaeophyceae) of seaweeds from specific geographic regions, for example DNA barcoding surveys from the Taiping Sea (Chen *et al*. 2022); northern Madagascar (Vieira *et al*. 2021); Bergen, Norway (Bringloe *et al*. 2019); Nome and the Beaufort Sea, Alaska (Bringloe *et al*. 2017, Bringloe & Saunders 2019) and Churchill, Canada (Saunders & McDevit 2010). Some others have focused only on the Rhodophyta of an island or coastal nation or region. For example, DNA barcoding surveys of the Rhodophyta have been carried out for Hawaii (Sherwood *et al*. 2010), Tristan da Cunha (Saunders *et al*. 2019) and Tunisia (Mangshisi *et al*. 2019). The results of all these studies showed that the true diversity of seaweeds had often been previously underestimated. Vieira *et al*. (2021) in their DNA barcoding survey of Madagascar, which represents an understudied area, more than doubled the known species from the region. Similar surveys have revealed cryptic alien species that would have otherwise gone undetected (Manghisi *et al*. 2019) and have also challenged the notion of wide distributions of certain species (Puckree-Padua *et al*. 2021).

The suite of DNA markers used for barcoding differs among the Rhodophyta, Chlorophyta and Phaeophycean seaweeds due to their diverse and complex evolutionary history (Rindi *et al*. 2011). This might explain why DNA barcoding surveys in the past tended to focus on a particular group of seaweeds and why many of them have focused on the Rhodophyta which account for a large majority of the global seaweed flora (Guiry & Guiry 2023). The Rhodophyta are routinely studied using the nuclear large subunit ribosomal RNA (rRNA) (LSU, ∼2 700 bp), the plastid marker ribulose-1,5-bisphosphate carboxylase/oxygenase large subunit (rbcL, ∼1 350 bp) and the protein-coding mitochondrial marker, the cytochrome c oxidase subunit I (COI∼1 232 bp) (Saunders 2005; Freshwater *et al*. 2010; Saunders & Moore 2013). Although each gene has its own set of advantages and limitations, the combination of different genes is often sufficient to overcome the challenges mentioned earlier (Saunders & Moore 2013).

It is important to remember that even though DNA barcoding was not initially intended for species discovery, it has been incredibly useful in this regard. Most DNA barcoding studies, however, are carried out in a conventional way, i.e. for species identification and comparison against a reference library. However, to date no DNA barcoding survey, at least for seaweeds, has been carried out in a taxonomically informed way in order to generate a well-curated reference library initially. Although the term Molecular Assisted Alpha Taxonomy (MAAT), coined by Saunders (2005), describes the utility of DNA barcoding for taxonomic research, it uses DNA barcodes from conventional barcoding surveys to guide further taxonomic work and morphological analyses (Cianciola *et al*. 2010). We argue that targeting geographic regions from which many many species have been described or where atypical or geographically isolated taxa are known to occur could similarly be useful in way that unites DNA barcoding and taxonomy. Despite the sequence of events (shot gun or target approaches), DNA barcoding, coupled with phylogenetic analyses, is a powerful tool and has challenged several well-established morphological species concepts (Savoie & Saunders 2019) and refined the classification of many taxa (eg. Nauer *et al*. 2016). Beyond its application in taxonomy, DNA barcoding has also been indispensable for biosecurity, wildlife forensics and certification fraud (Lowenstein *et al*. 2010) and conservation.

Approximately 850 species of seaweeds are known from South Africa (Bolton & Stegenga 2002), making it among the richer floras in the world (Lüning 1990) with high levels of endemism (Bolton 1994; Bolton & Stegenga 2002). South African collections feature in early works of seaweed taxonomy with many of the early plant collectors from the 1700s also collecting seaweeds, which were described in Europe by major seaweed taxonomists of the 18^th^ and 19^th^ centuries (Bolton 1999). South Africa thus has a relatively high level of endemic species and types, due to its geographic isolation, evolutionarily dynamic environment and early collecting history.

The South African continental coastline extends for almost 3000 kms and straddles two oceans influenced by contrasting current regimes (Griffiths *et al*. 2010; Smit *et al*. 2017). The warm Agulhas current flows south along the east and south coast whereas the cold Benguela current flows north along the west coast into Namibia and southern Angola (Griffiths *et al*. 2010). The environmentally dynamic west coast is further punctuated with wind-driven upwelling cells (Lutjeharms *et al*. 2001). The South African coast, together with Nambia, forms the temperate southern African marine realm which is further divided into three marine provinces and five ecoregions (Spalding *et al*. 2007). Although not included in Spalding’s delineation, southern Mozambique is geographically situated within southern Africa and the ranges of many species traverse this pollical boundary. The Agulhas Marine Province is confined within the political boundaries of South Africa while the Benguela Marine Province is shared with Namibia and southern Angola. Local biogeographic patterns in the South Africa seaweed flora correspond well with Spalding’s ecoregions (Spalding *et al*. 2007) and have been regarded by local phycologists as the cool temperate Benguela Marine Province (BMP) along the west coast, the warm temperate Agulhas Marine Province (AMP) along the south coast and the Indo-West Pacific Marine Province (IWP-MP) along the east coast (Anderson *et al*. 2009). The BMP and AMP are characterised by temperate seaweeds while the Indo-West Pacific Marine Province represents a wider tropical flora with northern KwaZulu-Natal marking the start of a more typical tropical seaweed flora (Bolton *et al*. 2004; Anderson *et al*. 2009). The south coast of South Africa, which corresponds to the AMP, holds the highest seaweed diversity with a peak in diversity around Port Alfred (coastal section 39 in Bolton & Stegenga 2002). Port Alfred is on a region of the coast influenced by a temporary upwelling cell (Lutjeharms *et al*. 2001) which creates an even more dynamic environment and as a result supports high levels of seaweed diversity (Anderson *et al*. 2009). Incidentally, Port Alfred has been a popular seaweed collecting site (often known as ‘the Kowie’ after the river which arises in Port Alfred, Bolton 1999) since the 1890s with Becker extensively collecting around the region, followed by Pocock’s collection which has contributed to a large collection of seaweeds housed at GRA. It is also where many seaweeds have their type localities. Anderson *et al*. (2009) identified the region as a high conservation priority because of its high seaweed diversity and endemism, but to date it has not been included in any MPA.

The diversity and distribution of South African seaweeds are relatively well documented (Stegenga *et al*. 1997; De Clerck *et al*. 2005; Anderson *et al*. 2016), and therefore provide a good background for DNA barcoding. We focused our collection efforts on the Port Alfred region as it is known to support high levels of diversity characteristic of the AMP and is where many seaweed species have been first described from South Africa. It is also home to a few monospecific genera. This study forms part of a preliminary investigation with a broader eventual aim to barcode the entire seaweed flora of South Africa. It also demonstrates the value of taxonomically guided DNA reference libraries that can be adopted for other marine organisms and in other regions of the world.

## Materials and methods

Several localities, Kenton-On-Sea (-33.6829; 26.6717), Port Alfred (-33.5979; 26.9017), Three Sisters (-33.5906; 26.8910) and the Great Fish River (-33.4274; 27.0993), around coastal sections 38 & 39, following Stegenga & Bolton (2002) covering *ca*. 50 km, in South Africa were sampled in different seasons during 2017-2018 (see Table A1 for more details). Samples were collected under the permits (reference numbers RES2017/07 and RES2018/100) issued by the Department of Environment, Forestry, and Fisheries (DEFF). A total of 158 specimens was collected during various sampling campaigns from the intertidal and near subtidal during spring low tides. Fresh specimens were assigned to morphological species (field IDs) using available taxonomic literature or field guides and in some cases additional anatomical work was carried out later to narrow the morphological identities. Specimens of interest (typically those representing new species) will be deposited in the Bolus Herbarium (BOL). All other voucher specimens are stored at the Seaweed Biorepository at the University of Cape Town. Specimens were assigned a unique identifier (D-number) which linked all samples to their corresponding metadata. Subsamples were preserved in silica-gel for DNA analyses and in some cases in 5% formalin/seawater for additional anatomical assessment.

Total genomic DNA was extracted from approximately 50 mg of silica dried tissue using a modified CTAB method (Clarke 2009). Dried tissue samples were homogenised into a fine powder using a TissueLyser (Mixer mill MM400, Retsch) prior to extraction with a CTAB buffer. In addition to the extraction buffer, 20 mg/ml Proteinase K was added to each sample and incubated overnight at 65°C. Pelleted DNA was dissolved in 50 μl MilliQ water and stored at room temperature before storage at -20°C until further analysis. The quality and quantity of the extracted DNA was assessed using a Nanodrop 2000/2000c spectrophotometer (Thermo Scientific). When necessary, RNase was added to samples. Samples were diluted as needed before PCR amplification.

Three gene regions were targeted for PCR amplification, (1) nuclear marker, LSU D2/D3, (2) mitochondrial marker, COI-5P and, (3) plastid marker, *rbc*L-3P using primers from Saunders and Moore (2013). Individual reactions were amplified in a total volume of 12.5 μl, which included 11.5 μl of PCR reaction mix and 1 μl of DNA template. The PCR reaction mix comprised of 25 mM MgCl2, 10 mM (2.5 mM each nucleotide) deoxynucleotide triphosphates (dNTPs), 10 x Taq reaction buffer, 10 μM of each primer and 5U Taq DNA polymerase. All amplifications included a negative control. PCR amplification profiles followed Saunders and McDevit (2012) and Reddy *et al*. (2018) and amplicons were verified on 1.5% (w/v) agarose gels using electrophoresis. The resultant bands were compared to a 1 Kb HyperLadder (Bioline) using the expected sizes per gene region as follows: 750 bp (LSU D2/D3), 664 bp (COI-5P) and 785 bp (rbcL-3P). Primer combinations for sequencing followed Saunders and Moore (2013) and bi-directional sequencing reactions were carried out using an ABI PRISM BigDye™ Terminator version 3.1 Cycle sequencing kit (Applied Biosystems). Sequencing reactions contained a final volume of 10 μl and composed of 5x BigDye Sequencing Buffer, 1 μM primer (forward or reverse) and 30 ng template DNA. The thermal profile for the sequencing reactions was as follows: denaturation at 94°C for 5 minutes, 35 cycles of 94°C for 10 seconds, 55°C for 10 seconds followed by 60°C for 4 minutes. Sequencing was carried out at the Central Analytical Facilities (CAF) at Stellenbosch University.

Sequence chromatographs were manually checked for ambiguities in BioEdit version 7.0.5.3 (Hall, 1999) and homologies were checked on BOLD: The Barcode of Life Data Systems (www.barcodinglife.org) and Basic Local Alignment Search Tool (BLAST). The identities of species were confirmed when the intraspecific diversity was lower than the interspecific diversity for known available sequences or using conventions from the literature (see Freshwater *et al*. 2010) or percentage cut-offs following Saunders *et al*. (2019).

In some instances where taxonomic anomalies were flagged, preliminary phylogenetic analyses were carried out using MrBayes version 3.2.6 (Huelsenbeck & Ronquist 2001) and the webserver RaXml (Stamatakis *et al*. 2008). Genetic distance comparisons with close relatives were compared using MEGA version X (Kumar *et al*. 2018). Details of the phylogenetic tree construction and genetic distance comparisons followed Reddy *et al*. (2020a) using concatenated multiple alignments when possible. The preliminary phylogenetic results are not presented here but were rather used to indicate which taxa require further taxonomic work including morphological assessment which was not within the scope of the present study.

In addition, species pages were compiled for all studied species according to guidelines by the South African National Biodiversity Institute (SANBI) and the Foundational Biodiversity Information Programme (FBIP).

## Results

A total of 220 DNA barcodes from three unlinked gene regions were generated, 85 for the LSU D2/D3 region which ranged in size from 232-364 bp, 76 for the *rbc*L-3P region ranging from 495-590 and 59 for the COI-5P region ranging from 340-504 bp. This translated into barcodes for 88 species and one species variety (Fig 1). The 88 studied species represented 13 orders, 24 families, 16 tribes and 55 genera.

**Figure 1.**
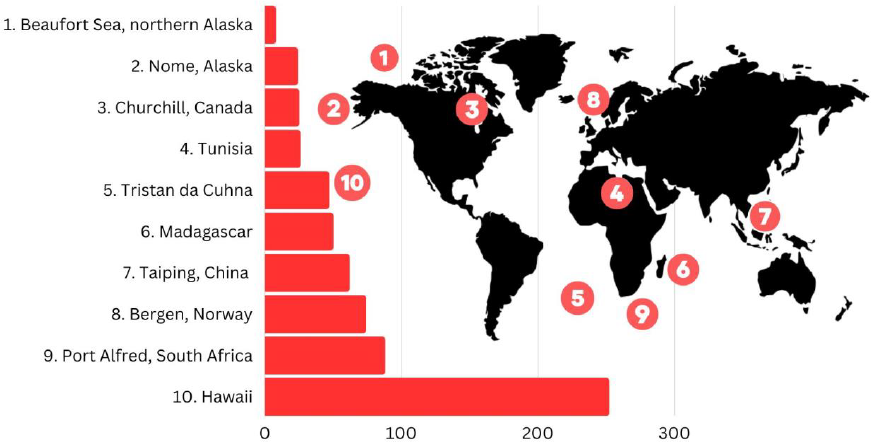
: A comparison of species represented by barcodes in studies from different regions in the world

Overall, new barcodes were generated for 73 species (62 species for LSU D2/D3; 35 species for COI-5P; 30 species for *rbc*L-3P), a little less than half (31) of which were species endemic to South Africa and an additional five species that were endemic to Southern Africa (Namibia, South Africa, Mozambique). In addition, 18 species (2 species for LSU D2/D3; 13 species for COI-5P; 4 species for *rbc*L-3P) that were previously represented by barcodes from either the BMP (Western Cape coast) or IWP-MP (KwaZulu-Natal coast) are now represented by barcodes from the AMP (Eastern Cape coast).

Our study also provided barcodes for nine species from their type locality (Port Alfred), and an additional eight species from near their type locality (AMP/Eastern Cape region). Only eight non-indigenous species (type localities outside South Africa) were confirmed to occur in South Africa and an additional six species presumed to be non-indigenous based on morphology were found to represent potentially new species as discussed below.

## Discussion

The long and often complicated process of alpha taxonomy and a lack of expertise on certain groups of marine organisms such as seaweeds (De Clerck *et al*. 2013) or knowledge of understudied regions of the world (Saunders *et al*. 2019; Vieira *et al*. 2021) make biodiversity studies all the more challenging (Reddy *et al*. 2021). While DNA barcoding is no substitute for alpha taxonomy (DeSalle 2006), it is certainly useful for understanding the overall biodiversity of a region and can be used to guide further research efforts (Reddy *et al*. 2020b; Reddy *et al*. 2023). The need for taxonomically guided DNA reference libraries however is much needed and is becoming more pertinent than ever, especially in an age of metabarcoding or eDNA (Cristescu and Hebert, 2018). While these high throughput sequencing methods developed for use by specialists and non-specialists alike show promise for future biodiversity assessments and monitoring, they first require well curated and taxonomically informed DNA reference libraries for them to be effective. DNA barcoding efforts vary from region to region or among taxa so these techniques will work better for some regions or groups of organisms than others. Furthermore, attaching a Linnean name to DNA barcodes should be carried out with caution as misidentifications may result in incorrect assumptions of diversity and distribution. Taxonomically guided DNA reference libraries are therefore critically needed as a baseline for future biodiversity, ecological or chemical studies.

In this study we initiate the construction of a taxonomically informed DNA reference library together with experts in the field for seaweeds in South Africa with the aim of continually adding to this baseline in the future. Similar efforts from other regions in the world or for other groups of marine organisms are encouraged. Our barcoding survey targeted a region (coastal sections 38-39, Anderson et al. 2009) along the coast of South Africa known to harbour some of the highest levels of red algal diversity and where many species have their type locality. Our studied species represented 21% of the diversity of the AMP even though our sampling campaign did not always include the collection of epiphytes or fragments of species which could have increased our estimate, not to mention the small portion of samples that we could not generate barcodes for. Our results support the hypothesis that Port Alfred can act as a surrogate of the seaweed diversity for the AMP. Preliminary phylogenetic analyses and comparison with the literature suggest that our dataset contained two potentially new genera (1 previously identified-‘*Tenebris’ sensu* Johnson 2017), 19 unidentified and potentially new species, one new species (Reddy *et al*. 2020b), one reinstated species (Reddy *et al*. 2023) and one new distribution record (this study). This indicates that almost a quarter of all species with barcodes in the present study represented novelty. *Ceramium arenarium* Simons, *Hypnea rosea* Papenfuss, *Tayloriella tenebrosa* (Harvey) Kylin, *Polyzonia elegans* Suhr, *Papenfussia laciniata* (Harvey) Kylin, *Pachychaeta brachyarthra* (Kützing) Trevisan, *Nienburgia serrata* (Suhr) Papenfuss, *Griffithsia confervoides* Suhr, *Bornetia repens* Stegenga, *Scinaia capensis* (Setchell) Huisman, *Botryocladia madagascariensis* G.Feldmann, *Heringia mirabilis* (C.Agardh) J.Agardh, *Arthrocardia corymbosa* (Lamarck) Decaisne, and *Gymnogongrus tetrasporifer* Papenfuss were among the studied species that were flagged for further taxonomic work including morphological assessment. In some instances, additional taxonomic work has already been carried out in separate publications and resulted in the description of one of new species, *C. africanum* Stegenga, M.M. Reddy, R.J. Anderson, Bolton (Reddy *et al*. 2020) and the reinstatement of *Plocamium robertiae* F.Schmitz ex Mazza, a species previously placed in synonymy with *P. corallorhiza* (Turner) Hooker f. & Harvey (Reddy *et al*. 2023).

When compared to similar barcoding studies carried out elsewhere in the world (eg. Bringloe *et al*. 2019; Saunders *et al*. 2019; Vieira *et al*. 2021), the number of species represented by barcodes in our relatively short DNA barcoding survey was remarkable. Our study produced barcodes for the second highest number of species from a comparison of 10 similar studies. It was only exceeded by a DNA barcoding survey by Sherwood *et al*. (2010) based on collections efforts from over a decade and covering a much wider geographic range than our study. It begs the question, what might a more comprehensive DNA barcoding survey carried out over a longer period of time in South Africa produce? The comparison of similar DNA barcoding surveys of the Rhodophyta around the world is not intended as a direct comparison of the seaweed floras in different regions as the sampling effort (collection trips and geographic area covered) as well as expertise will differ in different studies. It is merely intended to show the potential of similar DNA barcoding surveys of seaweeds in South Africa.

### Phylogenetic implications

Our DNA barcodes covered a wide range of phylogenetic diversity at different taxonomic levels. Eight of our studied species represented generitypes, a species designated to represent the type of a genus: *Papenfussia laciniata, Placophora binderi* (J.Agardh) J.Agardh, *Tayloriella tenebrosa, Pterosiphonia cloiophylla* (C.Agardh) Falkenberg, *Portieria hornemannii* (Lyngbye) P.C.Silva, *Polyzonia elegans, Codiophyllum natalense* J.E.Gray, *Heringia mirabilis*. The placement of the generitype anchors the taxonomic position of the genus in phylogenetic analyses and may result in systematic revisions and nomenclatural changes (Savoie & Saunders 2016; Savoie & Saunders 2019). For example, sequencing of topotype material of *P. cloiophylla* from South Africa saw the transfer of many other species previously assigned to *Pterosiphonia* to newly described or resurrected genera (Savoie & Saunders 2016). Similarly, the generitype *Carradoriella virgata* (C.Agardh) P.C.Silva (as *Polysiphonia virgata*) was used to resurrect and define *Carradoriella* (Savoie & Saunders 2019). Two genera in our study are monospecific, *Polyzonia* and *Heringia* and two others contain only three and two species each, *Placophora* and *Papenfussia*, respectively, and most are from South Africa. DNA barcodes for these local taxa generally confirmed their placement at the ordinal level but not always at the family level. For example, *H. mirabilis* was not resolved within the family Caulacanthaceae in which it is currently placed. Another example involved *Tayloriella* which includes four species, two of which have been sequenced but are distantly related to one another. However, as the generitype is designated as *T. tenebrosa*, the other sequenced species, *Tayloriella dictyurus* (J.Agardh) Kylin, will likely encounter systematic revision and name changes in the future, if the barcode truly represents the species in question. DNA barcodes for the remaining species are therefore needed to determine whether *Tayloriella* represents yet another monospecific genus from Southern Africa. The seaweed flora of warm temperate South Africa has a long evolutionary history (Hoek 1984) and unlike the northern hemisphere would have not encountered species fluctuations in response to ice ages (Wiencke *et al*. 1994). It is therefore likely that the long isolation of some taxa over time (Hommersand 1986) would have created distinct entities that warrant their own genera which is seen for other taxa mentioned earlier.

### Topotype barcodes and distribution implications

The notion of geographically widespread seaweed species is increasingly being challenged with the application of DNA. In at least two instances, non-indigenous species collected from South Africa (*Prionitis filiformis* Kylin; Type=USA or *Plocamium microcladioides* G.R.South & N.M.Adams; Type=NZ—see Reddy et al. 2023) were shown to be misidentifications. For example, *Prionitis filiformis* is distributed mostly around America with an unusual distribution record in South Africa. DNA barcodes generated in this study formed a separate clade from specimens of *Prionitis filiformis* from its type locality and therefore specimens from South Africa probably represent a new species and a new genus. However, a more comprehensive revision of the Halymeniales and stabilisation of the taxonomy of genera are needed before any species and genera are described. In the present study, we similarly identified, on morphological grounds, five other species with wide distribution ranges that had been previously recorded in South Africa based on morphology. These included, *Apoglossum ruscifolium* (Turner) J.Agardh (Type locality=Spain), *Caulacanthus ustulatus* (Turner) Kützing (Type locality=England), *Centroceras clavulatum* (C.Agardh) Montagne (Type locality=Peru), and *Botryocladia madagascariensis* (Type locality= Madagascar) and *Vertebrata foetidissima* (Cocks ex Bornet) Díaz-Tapia & Maggs (Type locality= England). However, in all cases our DNA results show that these names were misapplied and instead concealed potentially new species from the region. In fact, none of the six non-indigenous species (+ *Prionitis filiformis* (Type locality=California, USA)) recorded based on morphology have been confirmed with DNA in South Africa, therefore their presence in the region is highly doubtful. However, the presence of at least eight species with type localities outside South Africa, eg. *Chondracanthus acicularis* (Roth) Fredericq and *Antithamnionella spirographidis* (Schiffner) E.M.Wollaston were confirmed in the region. Although this makes up a relatively small percentage of our studied species, it reiterates the value of DNA confirmation for biodiversity assessments and accurate species estimates, as some species may be truly widely distributed while others may not.

Our study provided novel barcodes for 36 species endemic to Southern Africa. Barcodes produced for three endemic species, *Plocamium beckeri* F.Schmitz ex Simons, *Gracilaria denticulata* F.Schmitz ex Mazza, *Gigartina paxillata* Papenfuss for the first time were identical to sequences on GenBank going by a different species name (*P. beckeri*= *P. microcladioides*; COI-5P), (*G. paxillata* = *Gigartina polycarpa* (Kützing) Setchell & N.L.Gardner; COI-5P) and (*G. denticulata*= *Gracilaria huangii* S.-M.Lin & De Clerck; rbcl-3P). The misapplication of the name *P. microcladioides* for South African specimens of *Plocamium beckeri* F.Schmitz ex Simons has already been discussed (see Reddy *et al*. 2023). Similarly, *G. paxillata* is supported in this study as distinct from other members in the genus. The species however was not included in a revision of the *Gigartina* by Hommersand (1993) but has been mentioned in several local publications. Levitt (1998) suggested that the species was correctly placed in the genus *Gigartina* based on reproductive characters and his findings are supported here with DNA. Our specimen of *G. paxillata* was molecularly identical to another specimen collected from Port Alfred but identified as *Gigartina polycarpa* (Kützing) Setchell & N.L.Gardner by Hommersand and Fredericq (2003). These specimens were recovered in a well-supported sister clade with respect to specimens of *G. polycarpa* collected from the west coast of South Africa. *Gigartina polycarpa* is a species with a typical west coast distribution but has also been recorded at Port Alfred (Anderson *et al*. 2016). The specimen on GenBank (AF385664) however is clearly a misidentification of *G. paxillata*. Lastly, a sequence mislabelled as *G. huangii* S.-M.Lin & De Clerck (DQ296121) on GenBank was collected from South Africa and is correctly listed as *G. denticulata* F.Schmitz ex Mazza in Lin & De Clerck (2006). *Gracilaria huangii* (AY737438-40) is indeed distinct from *G. denticulata* but does not occur in South Africa. These examples, reiterate the value of generating barcodes for endemic species and combining it with phylogenetic analyses. For many other endemic species new barcodes were added. For example, 18s and *rbc*L barcodes were already available for *Gracilaria aculeata* and we add two new barcodes (LSU & COI) for the species. This is important because species are commonly represented by an uneven set of barcodes in public repositories so providing additional barcodes, particularly for regions that have an extensive reference collection, such as *rbc*L and COI will allow for a wider comparison of taxa. However, in some other cases, species were not represented by any barcodes until now, for example, *G. paxillata, G. tetrasporifer* Papenfuss, *P. lacinata, P. brachyarthra, Dasya echinata* Stegenga, Bolton & R.J.Anderson. In many of those cases their current taxonomic placement has not been supported and they were flagged for taxonomic revision (*P. lacinata, P. brachyarthra, D. echinata*).

The misidentification of specimens at both local and global scales not only influences the biodiversity estimates of species in a region but, in some cases, it also impacts the accepted distribution ranges of species. In our study, *Stirkia fujiiana* (Barros-Barreto & Maggs) Barros-Barreto & Maggs was recorded in South Africa for the first time. The species has likely been encountered before but was probably misidentified as one of our local species based on morphology (see Anderson *et al*. 2016). Similar misidentifications of two other species based on morphology, *Callithamnion collabens* L.M.McIvor & Maggs and *Vertebrata foetidissima* also had implications for the local distributions of these species. *Callithamnion collabens* was previously recorded along the BMP and AMP. However, specimens along the AMP were shown to be a new species (Reddy *et al*. 2020b), thereby restricting the distribution range of *C. collabens* to the BMP. Similarly, in the present study, a specimen morphologically identified as *V. foetidissima* was identical to a sequence of *Vertebrata urbana* (Harvey) Kuntze. The publicly available sequence of *V. urbana* (Díaz-Tapia *et al*. 2017) was however collected from outside the known local distribution of the species (Stegenga *et al*. 1997). Sequences from the type locality of *V. urbana* are therefore needed to verify whether the range of the species extends to the the AMP (south coast). What is certain is that the presence of *Polysiphonia foetidissima* has not yet been confirmed in South Africa based on DNA. The ranges of one other species, *Laurencia dichotoma* Francis, Bolton, Mattio & R.J.Anderson (by ca. 1000 km) and one species variety *Laurencia pumila var dehoopiensis* C.Francis, J.J.Bolton, Mattio & R.J.Anderson (by ca. 700 km), are also significantly extended. Both were previously only known from their type localities.

The South African coast straddles two oceans with environmentally contrasting conditions (Smit *et al*. 2017). The verification of species along cool or warm temperate and tropical regions of the coastlines is therefore necessary to verify the identity of species with truly wide local distributions or to detect cases of cryptic speciation (Reddy 2018). In our study, we therefore noted and compared new barcodes for species generated from different Marine Provinces, the BMP (Western coast), IWP-MP (East coast) or AMP (South coast). In most cases, species with barcodes from different Marine Provinces along the coastline were generally resolved within the same species with little to no genetic divergence. However, in at two cases biogeographic structure related to Marine Provinces were observed within species: *P. suhrii* (1% *rbc*L) and *P. cornea* (0.7% *rbc*L) and warrant further investigation.

## Conclusion

The South African seaweed flora is relatively well studied from a morphological perspective (Stegenga *et al*. 1997; De Clerck *et al*. 2005; Anderson *et al*. 2016) and provided us with an opportunity to link Linnean names to DNA sequences in a taxonomically guided way. The impact of DNA has drastically improved the knowledge of our local seaweed flora, but much work remains in this regard as only 24% of the known flora is represented by barcodes (Reddy; unpublished data). In this study, our barcoding approach was slightly different from the conventional approach where DNA barcodes are compared to a reference library. Instead, we aimed to provide a taxonomically guided DNA reference library in the first place with emphasis on species endemic to Southern Africa and including species from their type locality and for monospecific or geographically isolated genera including several generitypes. This approach ensured that we not only provided barcodes but also avoided misapplication of names or mislabelling of taxa as is commonly encountered on public sequence repositories. Our morphological and field identifications were largely in agreement with existing DNA barcodes.

Overall, these results not only contribute novel barcodes for the South African seaweed flora but also provide a baseline for future seaweed work. For example, eDNA metabarcoding has been touted as a new tool for biodiversity monitoring, detecting non-invasive species and recording changes in distribution patterns over a temporal and spatial scale (Ruppert *et al*. 2021). The efficacy of eDNA however depends on a reliable reference library for comparisons. We envision that DNA barcoding reference libraries linked to voucher specimens (Reddy *et al*. 2021, this study) such as that provided in the present study should be the first step in documenting biodiversity, followed by metabarcoding and eDNA.

## Acknowledgements

We are grateful to Dave Dyer and Chris Boothroyd and Derek Kemp from the Seaweed Unit (DEEF/UCT) for assistance in the field. We also thank SeaKeys and the University of Cape Town URC (MMR, postdoc) for funding.

## Legends

**Figure 1:** a comparison of DNA barcoding surveys, comparing only the Rhodophyta, from different regions in the world (y-axis) with the number of species represented by barcodes (x-axis). The corresponding numbers show their approximate location on a schematic view of a world map.

**Table A1:** Specimen collection information, DNA barcodes and notes

**Table S1:** List of DNA barcoding surveys used to generate figure 1

## References

Anderson R.J., Bolton J.J. & Stegenga H. 2009. — Using the biogeographical distribution and diversity of seaweed species to test the efficacy of marine protected areas in the warm-temperate Agulhas Marine Province, South Africa. Diversity and Distributions 15: 1017–1027.

Anderson R.J., BOLTON J.J. & Stegenga, H. 2016. — Seaweeds of the South African South Coast. World-wideweb electronic publication, University of Cape Town. Retrieved fromhttp://southafrseaweeds.uct.ac.za, (Accessed 2018–2023).

Bolton J.J., Leliaert F., Declerck O., Anderson R.J., Stegenga H., Engledow H.E. & Coppejans, E. 2004. — Where is the western limit of the tropical Indian Ocean seaweed flora? An analysis of intertidal seaweed biogeography on the east coast of South Africa. Marine Biology 144: 51–59.

Bolton J. J. 1994 — Global seaweed diversity: patterns and anomalies. Botanica Marina 37: 241–245.

Bolton J.J. 1999. Seaweed systematics and diversity in South Africa: an historical account. Transactions of the Royal Society of South Africa 54: 167–177.

Bolton J.J. & Stegenga H. 2002. — Seaweed species diversity in South Africa. South African Journal of Marine Science 24: 9–18.

Bringloe T.T., SjØTun K. & Saunders G.W. 2019. — A DNA barcode survey of marine macroalgae from Bergen (Norway). Marine Biology Research 15: 580–589.

Chapman, R.L. 2013. — Algae: the world’s most important “plants” an introduction. Mitigation and Adaptation Strategies for Global Change 18: 5–12.

Clarke, J.D. 2009. — Cetyltrimethyl ammonium bromide (CTAB) DNA miniprep for plant DNA isolation. Cold Spring Harbor Protocols. 4: 5177–5179.

Cristescu, M.E. and Hebert, P.D. 2018. Uses and misuses of environmental DNA in biodiversity science and conservation. Annual Review of Ecology, Evolution, and Systematics 49: 209–230.

De Clerck O., Guiry M.D., Leliaert F., Samyn Y. & Verbruggen H. 2013. — Algal taxonomy: a road to nowhere?. Journal of Phycology 49: 215–225.

Desalle R. 2006. — Species discovery versus species identification in DNA barcoding efforts: Response to Rubinoff. Conservation Biology 20: 1545–1547.

De Clerck O., Bolton J.J., Anderson R.J., Coppejans E., Bolton J. & Anderson R.L. 2005. — Guide to the seaweeds of KwaZulu-Natal. Scripta Botanica Belgica, 33. Flanders Marine Institute (VLIZ)/Flemish Community/National Botanic Garden of Belgium: Meise. ISBN 90-72619-64-1. 294, ill. pp.

DíAz-Tapia, P., Mcivor, L., Freshwater, D.W., Verbruggen, H., Wynne, M.J. & Maggs, C.A. 2017. The genera Melanothamnus Bornet & Falkenberg and Vertebrata SF Gray constitute welldefined clades of the red algal tribe Polysiphonieae (Rhodomelaceae, Ceramiales). European Journal of Phycology, 52: 1–30.

DíAz-Tapia P., Ly M. & Verbruggen H. 2020. — Extensive cryptic diversity in the widely distributed Polysiphonia scopulorum (Rhodomelaceae, Rhodophyta): molecular species delimitation and morphometric analyses. Molecular Phylogenetics and Evolution 152: 106909.

Cianciola E.N., Popolizio T.R., Schneider C.W. & Lane C.E. 2010. — Using molecular-assisted alpha taxonomy to better understand red algal biodiversity in Bermuda. Diversity 2: 946–958.

Freshwater D.W., Tudor K., O’Shaughnessy, K. & Wysor B. 2010. — DNA barcoding in the red algal order Gelidiales: Comparison of COI with rbcL and verification of the “barcoding gap”. Cryptogamie Algologie 31: 435–449.

Griffiths C.L., Robinson T.B., Lange L. & Mead A. 2010. — Marine biodiversity in South Africa: An evaluation of current states of knowledge. PLoS One 5: 1–13.

Guiry M.D. & Guiry G.M. 2023 — AlgaeBase. World-wide electronic publication, National University of Ireland, Galway. Retrieved from:http://www.algaebase.org, (Accessed 2020-2023).

Hall T. 1999. — BioEdit: user-friendly biological sequence alignment editor and analysis program for Windows 95/98/NT. Nucleic Acids Symposium Series 41: 95–98.

Hebert P.D.N., Cywinska A., Ball S.L. & Dewaard J.R. 2003a. — Biological identifications through DNA barcodes. Proceedings of the Royal Society B: Biological Sciences 270: 313–321.

Hebert P.D.N., Ratnasingham S. & Dewaard J.R. 2003b. — Barcoding animal life: cytochrome c oxidase subunit 1 divergence among closely related species. Proceedings of the Royal Society B: Biological Sciences 270: S96–S99.

Hommersand M.H. 1986. — The biogeography of the South African marine red algae. Botanica Marina 29: 257–270.

Hommersand M. H., Guiry M. D., Fredericq S. & Leister G. L. 1993. — New perspectives in the taxonomy of the Gigartinaceae (Gigartinales, Rhodophyta). Hydrobiologia 260: 105–120.

Hommersand M.H. & Fredericq S. 2003. — Biogeography of the red seaweeds of the South African west coast: a molecular approach. In Chapman, A.R.O., Anderson, R.J., Vreeland, V.J.and Davison, I.R. (eds) Proceedings of the 17th International Seaweed Symposium. Cape Town, Oxford University Press, pp. 325–336.

Johnson V.J. 2017. — Molecular systematics of the genus Hypnea (Rhodophyta) in South Africa, with the description of a new genus, Tenebris (Cystocloniaceae, Rhodophyta). MSc thesis, University of Cape Town, South Africa.

Kumar S., Stecher G., Li M., Knyaz C. & Tamura K. 2018. — MEGA X: Molecular evolutionary genetics analysis across computing platforms. Molecular Biology and Evolution 35: 1547–1549.

Levitt G 1998. — Studies on carrageenophytes of the Western Cape, South Africa: ecology, management and systematics. PhD Thesis, University of Cape Town, 183 pp.

Lin S.-M. & De Clerck O. 2006. — A new flattened species of Gracilaria (Gracilariales, Rhodophyta) from Taiwan. Cryptogamie Algologie 27: 233–244.

Ligtvoet A., Maier F., Sijtsma L., Van Den Broek L.A.M., Doranova A., Eaton D., Guznajeva T., Kals J., Le Gallou M., Poelman M. & Saes L. 2019. — Blue Bioeconomy Forum: Roadmap for the blue bioeconomy. European Commission.

Lindsey Zemke-White W. & Ohno M. 1999. — World seaweed utilisation: an end-of-century summary. Journal of applied Phycology 11: 369–376.

Lowenstein J.H., Burger J., Jeitner C.W., Amato G., Kolokotronis S.O. & Gochfeld M. 2010. — DNA barcodes reveal species-specific mercury levels in tuna sushi that pose a health risk to consumers. Biology Letters 6: 692–695.

LÜNing, K. 1990 — Seaweeds: their Environment, Biogeography and Ecophysiology. New York; Wiley-Interscience: 527 pp.

Lutjeharms J.R.E. & Bornman T.G. 2010. — The importance of the greater Agulhas Current is increasingly being recognised. South African Journal of Science 106: 1–4.

Manghisi A., Miladi R., Minicante S.A., Genovese G., Le Gall L., Abdelkafi S., Saunders G.W. & Morabito, M. 2019. — DNA barcoding sheds light on novel records in the Tunisian red algal flora. Cryptogamie Algologie 40: 5–27.

Miladi R., Manghisi A., Minicante S.A., Genovese G., Abdelkafi S. & Morabito M. 2018. — A DNA barcoding survey of Ulva (Chlorophyta) in Tunisia and Italy reveals the presence of the overlooked alien U. ohnoi. Cryptogamie Algologie 39: 85–107.

Moritz C. & Cicero C. 2004. — DNA barcoding: Promise and pitfalls. PLoS Biology 2: e354.

Nauer F., Cassano V. & Oliveira M.C. 2016. — Hypnea wynnei and Hypnea yokoyana (Cystocloniaceae, Rhodophyta), two new species revealed by a DNA barcoding survey on the Brazilian coast. Phytotaxa 268: 123–134.

Puckree-Padua C.A., Gabrielson P.W. & Maneveldt G.W. 2021. DNA sequencing reveals three new species of Chamberlainium (Corallinales, Rhodophyta) from South Africa, all formerly passing under Spongites yendoi. Botanica Marina 64: 19–40.

Reddy M.M. 2018. — Taxonomy and Systematics of the Bangiales (Rhodophyta) in South Africa Using an Integrative Approach. Doctoral thesis. University of Cape Town, Cape Town, South Africa.

Reddy M.M., Jennings L. & Thomas O.P. 2021. — Marine Biodiscovery in a Changing World. Progress in the Chemistry of Organic Natural Products 116: 1–36.

Reddy M.M., Clerck O.D., Leliaert F., Anderson R.J. & Bolton J.J. 2018. — A rosette by any other name: species diversity in the Bangiales (Rhodophyta) along the South African coast. European Journal of Phycology 53: 67–82.

Reddy M.M., De Clerck O., Leliaert F., Anderson R.J. & Bolton J.J. 2020a. — An appraisal of the genus Pyropia (Bangiales, Rhodophyta) from southern Africa based on a multi-gene phylogeny, morphology and ecology, including the description of Pyropia meridionalis sp. nov. South African Journal of Botany 131: 18–32.

Reddy M.M., Stegenga H., ANDERSON R.J. & Bolton J.J. 2020b. — An updated species inventory of Callithamnion sensu lato Rhodophyta, Callithamniaceae in South Africa with the description of Callithamnion africanum sp. nov. Phytotaxa 461: 139–154.

Ruppert K.M., Kline R.J. & Rahman M.S. 2019. — Past, present, and future perspectives of environmental DNA (eDNA) metabarcoding: A systematic review in methods, monitoring, and applications of global eDNA. Global Ecology and Conservation 17: p.e00547.

Reddy M.M., Du Plessis J., Roodt-Wilding R., Anderson R.J. & Bolton J.J. 2023. — The reinstatement of Plocamium robertiae (Rhodophyta, Plocamiales) and an updated species inventory of the genus in South Africa. Phycologia 62: 194–202.

Rindi F., Soler-Vila A. & Guiry M.D. 2011. — Taxonomy of marine macroalgae used as sources of bioactive compounds. In Hayes, M. (ed) Marine Bioactive Compounds: Sources, Characterization and Applications. Boston, Springer pp. 1–53.

Ronquist F. & Huelsenbeck J.P. 2003. — MrBayes 3: Bayesian phylogenetic inference under mixed models. Bioinformatics 19: 1572–1574.

Saunders G.W. 2005 — Applying DNA barcoding to red macroalgae: a preliminary appraisal holds promise for future applications. Proceedings of the Royal Society B: Biological Sciences 360: 1879–1888.

Saunders G.W. & Moore T.E. 2013. — Refinements for the amplification and sequencing of red algal DNA barcode and RedToL phylogenetic markers: a summary of current primers, profiles and strategies. Algae 28: 31–43.

Saunders G.W., Brooks C.M. & Scott S. 2019. — Preliminary DNA barcode report on the marine red algae (Rhodophyta) from the British Overseas Territory of Tristan da Cunha. Cryptogamie Algologie 40: 105–117.

Saunders G.W. & Mcdonald B. 2010. — DNA barcoding reveals multiple overlooked Australian species of the red algal order Rhodymeniales (Florideophyceae), with resurrection of Halopeltis J. Agardh and description of Pseudohalopeltis gen. nov. Botany 88: 639–667.

Saunders G.W. & Mcdevit D.C. 2012. Methods for DNA barcoding photosynthetic protists emphasizing the macroalgae and diatoms. Molecular Biology 858: 207–222.

Saunders G.W. & Mcdonald B., 2010. — DNA barcoding reveals multiple overlooked Australian species of the red algal order Rhodymeniales (Florideophyceae), with resurrection of Halopeltis J. Agardh and description of Pseudohalopeltis gen. nov. Botany 88: 639–667.

Savoie A.M. & Saunders G.W. 2016. — A molecular phylogenetic and DNA barcode assessment of the tribe Pterosiphonieae (Ceramiales, Rhodophyta) emphasizing the Northeast Pacific. Botany 94: 917–939.

Savoie A.M. & Saunders G.W. 2019. — A molecular assessment of species diversity and generic boundaries in the red algal tribes Polysiphonieae and Streblocladieae (Rhodomelaceae, Rhodophyta) in Canada. European Journal of Phycology 54: 1–25.

Seagrief S.C. 1988. — Marine algae. A Field Guide to the Eastern Cape Coast. Wildlife Society of Southern Africa, Grahamstown, pp.35–72.

Sherwood A.R., Kurihara A., Conklin K.Y., Sauvage T. & Presting G.G. — 2010. The Hawaiian Rhodophyta Biodiversity Survey (2006-2010): a summary of principal findings. BMC Plant Biology 10: 1–29.

Steinhagen S., Karez R. & Weinberger, F. 2019. — Surveying seaweeds from the Ulvales and Fucales in the world’s most frequently used artificial waterway, the Kiel Canal. Botanica Marina 62: 51–61.

Stamatakis A., Hoover P. & Rougemont J. 2008. — A rapid bootstrap algorithm for the RAxML Web servers. Systematic Biology 57: 758–771.

Spalding M.D., Fox H.E., Allen G.R., Davidson N., FerdaÑA Z.A., Finlayson M., Halpern B.S., Jorge M.A., Lombana A., Lourie S.A., Martin K.D., Mcmanus E., Molnar J., Recchia C.A. & Robertson J. 2007. — Marine ecoregions of the world: A bioregionalization of coastal and shelf areas. BioScience 57: 573–583.

Smit, A.J., Bolton, J.J. & Anderson, R.J. 2017 — Seaweeds in two oceans: Beta-diversity. Frontiers in Marine Science 4: 1–16.

Stegenga H., Bolton J.J. & Anderson, R.J. 1997. Seaweeds of the South African West Coast. Contributions from the Bolus Herbarium 18: 1–655.

Vieira C., De RAMON N’Yeurt A., Rasoamanendrika F.A., D’Hondt S., Tran L.A.T., Van Den Spiegel D., Kawai H. & De Clerck O. 2021. — Marine macroalgal biodiversity of northern Madagascar: morpho-genetic systematics and implications of anthropic impacts for conservation. Biodiversity and Conservation 30: 1501–1546.

Wiencke C., Bartsch I., Bischoff B., Peters A.F. & Breeman A.M. 1994. — Temperature requirements and biogeography of Antarctic, Arctic and amphiequatorial seaweeds. Botanica Marina 37: 247–259.

Wilson J.-J., Sing K.-W. & Jaturas N. 2018. — DNA barcoding: Bioinformatics workflows for beginners. Encyclopedia of Bioinformatics and Computational Biology 2012: 985–995.

