## Supplementary material for "A taxonomically informed DNA reference library to facilitate future biodiversity assessments and monitoring: a case study using seaweeds along a tropical-temperate transition zone in South Africa": Table A1

TableA1:

| **Taxon** | **Type locality** Site, Marine Province, Country | **ID** | **Date** | **Site** | **LSU; *rbc*L; COI;** | **Notes** |
|---|---|---|---|---|---|---|
| **Bangiales** | | | | | | |
| Bangiaceae | | | | | | |
| *Porphyra agulhensis* | Algoa Bay, AMP, South Africa | D2775 | 06-10-2017 | Kenton-on-Sea | 1; 1; 1; | Described in Reddy 2018 |
| **Bonnemaisoniales** | | | | | | |
| Bonnemaisoniaceae | | | | | | |
| *Delisea flaccida* (Suhr) Papenfuss | Algoa Bay, AMP, South Africa | D2777 | 06-10-2017 | Kenton-on-Sea | **1**; 1; ND; | New barcode for SA endemic from near its type locality. Clusters with other species in the genus, although the type species has not yet been sequenced |
| **Ceramiales** | | | | | | |
| Ceramiales sp. 1 | - | D2805; D2806 | 06-10-2017 | Kenton-on-Sea | ND; **1**; **2**; | Closest relatives (*rbc*L -89%) and (COI-87%) include other members of the Ceramiales. Potentially new species |
| Ceramiales sp. 2 | - | D2857 | 09-10-2017 | Kenton-on-Sea | ND; ND; **1**; | Closest relatives (COI-87%) include other members of the Ceramiales. This species was morphologically identified as *C. arenarium*. Potentially new species |
| Delesseriaceae | | | | | | |
| Phycodryeae | | | | | | |
| *Nienburgia serrata* (Suhr) Papenfuss* | Algoa Bay, AMP, South Africa | D2908 | 29-05-2018 | Three Sisters | **1**; **1**; **1**; | First barcodes for the species resolve it within a clade containing the generitype for *Polyneura* and shows a distant relationship with the generitype of *Neinburgia* |
| Callithamniaceae | | | | | | |
| Callithamnieae | | | | | | |

| | | | | | | |
|---|---|---|---|---|---|---|
| *Callithamnion africanum* Stegenga, M.M Reddy, R.J.Anderson & Bolton | Platboom, AMP, South Africa | D2801-2; D2852-3 | 06-10-2017 | Kenton-on-Sea | 4;<br>3;<br>4; | *Callithamnion africanum* (Reddy et al. 2020); misidentified as *C. collabens* which is likely restricted to the west coast of SA |
| *Callithamnion stuposum* Suhr* | Cape of Good Hope, BMP, South Africa | D2803 | 06-10-2017 | Kenton-on-Sea | 1;<br>1;<br>ND; | Not a member of the *Callithamnion* s.s. *Callithamnion* s.l. in need of taxonomic revision |
| Spyridieae<br>*Spyridia cupressina* Kützing | Cape of Good Hope, BMP, South Africa | D2778 | 06-10-2017 | Kenton-on-Sea | **1**;<br>1;<br>**1**; | Confirmed within the genus containing a barcode of the generitype but note large genetic variably (15% rbcL) within the genus |
| *Spyridia plumosa* F.Schmitz ex J.Agardh | Port Alfred, AMP, South Africa | D2823; D2925 | 07-10-2017 | Port Alfred | **2**;<br>ND;<br>ND; | New barcodes from type locality |
| Dohrnielleae<br>*Antithamnionella spirographidis* (Schiffner) E.M.Wollaston | Trieste, Italy | D2830 | 07-10-2017 | Port Alfred | **1**;<br>1;<br>**1**; | First ever COI barcode for *Antithamnionella spirographidis*. Confirmed (DNA) distribution in SA, Netherlands, USA |
| Ceramieae<br>*Centroceras* sp. 1 | - | D2850 | 09-10-2017 | Kenton-on-Sea | **1**;<br>ND;<br>**1**; | Potentially new species |
| *Centroceras* sp. 2 | - | D2919 | 29-05-2018 | Three Sisters | **1**;<br>**1**;<br>**1**; | Potentially new species |
| *Stirkia fujiiana* Barros-Barreto & Maggs | Espiriito Santo, Brazil | D2921 | 29-05-2018 | Great Fish River | ND;<br>1;<br>ND; | New distribution record |
| Delesseriaceae<br>*Dasya echinata* Stegenga, Bolton & R.J.Anderson* | Brandfontein, BMP, South Africa | D2842 | 09-10-2017 | Kenton-on-Sea | **1**;<br>ND;<br>ND; | First barcode for the species |

| | | | | | | |
|---|---|---|---|---|---|---|
| *Heterosiphonia arenaria* Kylin | Port Elizabeth, AMP, South Africa | D2848 | 09-10-2017 | Kenton-on-Sea | **1**; ND; ND; | First barcode for the species confirms its placement in the genus, closet relative *H. plumosa* (KJ961357.1) |
| *Heterosiphonia* sp. 1 | - | D2845 | 09-10-2017 | Kenton-on-Sea | ND; **1**; ND; | Potentially new species. We were unable to compare *H. arenaria* and this species because of varying gene regions amplified. However, the closet relative for each appears to be a different species of *Heterosiphonia* which suggests that they may represent different species |
| Papenfussieae | | | | | | |
| *Papenfussia laciniata* (Harvey) Kylin* | False Bay, BMP, South Africa | D2836 | 09-10-2017 | Kenton-on-Sea | **1**; 1; ND; | Closest relative (*rbc*L-90%) *Dasysiphonia* sp. 2WA (MW698706.1). Confirmed placement in the Ceramiales |
| Apoglosseae | | | | | | |
| *Paraglossum* sp. 1 | - | D2854 | 09-10-2017 | Kenton-on-Sea | ND; **1**; ND; | Potentially new species |
| Rhodomelaceae | | | | | | |
| *Chondria* sp. 1 | - | D2834; D2918 | 09-10-2017; 29-05-2018 | Kenton-on-Sea; Three Sisters | **2**; ND; ND; | Potentially new species |
| *Pachychaeta brachyarthra* (Kützing) Trevisan* | Cape Agulhas, BMP, South Africa | D2895 | 28-05-2018 | Port Alfred | **1**; 1; ND; | First barcodes for the species. *Pachychaeta griffithsioides* the type of the genus was synomized with this species |
| *Placophora binderi* (J.Agardh) J.Agardh | Cape of Good Hope, BMP, South Africa | D2829 | 07-10-2017 | Port Alfred | **1**; **1**; ND; | First barcode for *Placophora binderii* confirms its placement in the genus and family |
| *Tayloriella tenebrosa* (Harvey) Kylin* | False Bay, BMP, South Africa | D2847; D2942 | 09-10-2017; 30-05-2018 | Kenton-on-Sea; Great Fish River | ND; *2*; ND; | First barcodes for the species from the EC |
| Bostrychieae | | | | | | |

| | | | | | | |
|---|---|---|---|---|---|---|
| *Bostrychia intricata* (Bory) Montagne | Falkland Islands | D2809 | 06-10-2017 | Kenton-on-Sea | ND; ND; *1*; | First barcode for the species from the EC. Confirmed (DNA) distribution along the WC, EC and KZN in SA |
| Laurencieae | | | | | | |
| *Laurencia alfredensis* Francis, Bolton, Mattio & R.J.Anderson | Port Alfred, AMP, South Africa | D2898; D2904 | 28-05-2018; 28-05-2018 | Port Alfred; Port Alfred | ND; 2; **2**; | New barcodes from the type locality |
| *Laurencia corymbosa* J.Agardh | Cape of Good Hope, BMP, South Africa | D2787 | 06-10-2017 | Kenton-on-Sea | **1**; ND; **1**; | New barcodes for the species from SA |
| *Laurencia digitata* Francis, Bolton, Mattio & R.J.Anderson | Koppie Allen, AMP, South Africa | D2839 | 09-10-2017 | Kenton-on-Sea | 1; *1;* **1**; | Morphologically identified as *L. pumila* and resolved as *L. digitata* |
| *Laurencia flexuosa* Kützing | Cape of Good Hope, BMP, South Africa | D2786 | 06-10-2017 | Kenton-on-Sea | 1; *1;* 1; | |
| *Laurencia glomerata* (Kützing) Kützing | Cape of Good Hope, BMP, South Africa | D2789 | 06-10-2017 | Kenton-on-Sea | **1**; 1; **1**; | |
| *Laurencia griseaviolacea* M.J.Wynne | Clovelly, BMP, South Africa | D2788 | 06-10-2017 | Kenton-on-Sea | **1**; ND; **1**; | *Laurencia griseaviolacea* was validly published before the name *L. stegengae* was proposed by Francis et al. 2017. New barcodes for SA endemic |
| *Laurencia dichotoma* Francis, Bolton, Mattio & R.J.Anderson | Sodwana Bay, IWP-MP, South Africa | D2779 | 06-10-2017 | Kenton-on-Sea | 1; 1; ND; | Prior to this study *L. dichotoma* was only known from its type locality. Its distribution range is now extended to PA |
| *Laurencia pumila* (Grunow) Papenfuss | Durban, KZN | D2840 | 09-10-2017 | Kenton-on-Sea | **1**; ND; **1**; | New barcodes for Southern African endemic |
| *Laurencia pumila var dehoopiensis* Francis, Bolton, Mattio & R.J.Anderson | De Hoop, AMP, South Africa | D2825 | 07-10-2017 | Port Alfred | 1; 1; ND; | Prior to this study this species was only known from its type locality. Its range is now extended to PA. Specimen found growing epilithic as opposed to epiphytic as in previous collections |
| Polysiphonieae | | | | | | |

| | | | | | | |
|---|---|---|---|---|---|---|
| *Melanothamnus incomptus* (Harvey) Díaz-Tapia & Maggs | False Bay, BMP, South Africa | D2797;<br>D2862 | 06-10-2017 | Kenton-on-Sea | **1;**<br>**1;**<br>**1;** | First barcode for the species confirms its placement in *Melanothamus.* Southern African endemic |
| *Polysiphonia kowiensis* Stegenga, J.J. Bolton & R.J. Anderson | Port Alfred, AMP, South Africa | D2831 | 06-10-2017 | Kowie | ND;<br>ND;<br>**1;** | First barcode for the species suggests taxonomic revision |
| *Polysiphonia namibiensis* Stegenga & Engeldow | Hottentots Bay, Namibia | D2900 | 28-05-2018 | Port Alfred | **1;**<br>ND;<br>ND; | First barcode for the species suggests taxonomic revision. Southern African endemic |
| *Melanothamus* sp. 1 MMR2023 | - | D2906;<br>D2917 | 29-05-2018;<br>29-05-2018 | Three Sisters;<br>Three Sisters | **2;**<br>**2;**<br>**2;** | Potentially new species |
| *Vertebrata* sp. 1 MMR2023 | - | D2851 | 09-10-2017 | Kenton-on-Sea | **1;**<br>**1;**<br>**1;** | Morphologically identified as *Polysiphonia cf urbana* but probably a new species of *Vertebrata.* |
| *Melanothamus* sp. 2 MMR2023 | - | D2863 | 09-10-2017 | Kenton-on-Sea | **1;**<br>**1;**<br>**1;** | Potentially new species of *Melanothamus.* |
| *Vertebrata urbana* (Harvey) Kuntze | Table Bay, BMP, South Africa | D2864 | 09-10-2017 | Kenton-on-Sea | ND;<br>**1;**<br>ND; | Morphologically identified as *Polysiphonia foetidissima*. Barcode confirms its placement in *Vertebrata.* Southern African endemic |
| Polyzonieae | | | | | | |
| *Polyzonia elegans* Suhr | Algoa Bay, AMP, South Africa | D2902 | 28-05-2018 | Port Alfred | ND;<br>*1;*<br>ND; | First barcode from near the type locality of the species |
| Pterosiphonieae | | | | | | |
| *Pterosiphonia cloiophylla* (C.Agardh) Falkenberg | Cape of Good Hope, AMP, South Africa | D2907 | 29-05-2018 | Three Sisters | **1;**<br>*1;*<br>*1;* | New barcodes for the species from the EC |
| *Pterosiphonia stegengae* Savoie & G.W.Saunders | Stillbaai, BMP, South Africa | D2905 | 28-05-2018 | Port Alfred | **1;**<br>1;<br>1; | |
| Griffithsieae | | | | | | |

| | | | | | | |
|---|---|---|---|---|---|---|
| *Bornetia repens* Stegenga* | Hluleka, BMP, South Africa | D2849 | 09-10-2017 | Kenton-on-Sea | **1**;<br>ND;<br>ND; | First barcode generated for a species endemic to southern Africa resolved it within the Ceramiales |
| *Griffithsia confervoides* Suhr* | False Bay, BMP, South Africa | D2822 | 07-10-2017 | Port Alfred | ND;<br>**1**;<br>ND; | First barcode for a Southern African endemic species. Closely related to other species of *Griffithsia* but distant from the type of the genus |
| Wrangelieae<br>*Wrangelia purpurifera* J.Agardh | Cape of Good Hope, BMP, South Africa | D2920 | 29-05-2018 | Three Sisters | **1**;<br>**1**;<br>**1**; | First barcodes for the species confirm its placement in *Wrangelia* |
| Corallinales<br>Lithophyllaceae<br>Amphiroeae<br>*Amphiroa beauvoisii* J.V.Lamouroux | Portugal | D2785 | 06-10-2017 | Kenton-on-Sea | **1**;<br>**1**;<br>ND; | These represent the first LSU and *rbc*L barcodes for the species. Sequences from Tunisia under the same name are different from SA material |
| *Amphiroa capensis* Areschoug | False Bay, BMP, South Africa | D2780 | 06-10-2017 | Kenton-on-Sea | **1**;<br>ND;<br>ND; | First barcode for the species. This species was not included in Kogame et al. 2017 |
| Corallinaceae<br>*Arthrocardia corymbosa* (Lamarck) Decaisne* | Cape of Good Hope, BMP, South Africa | D2791 | 06-10-2017 | Kenton-on-Sea | **1**;<br>ND;<br>ND; | See generic replacement (Kogame et al. 2017) |
| *Jania cultrata* (Harvey) J.H.Kim, Guiry & H.-G.Choi | Durban, IWP-MP, South Africa | D2838 | 08-10-2017 | Kenton-on-Sea | **1**;<br>**1**;<br>1; | Morphologically identified in the field as *Jania sagittata* but resolved as *J. cultrate*. New barcodes from SA |
| *Corallina* sp. 1 | - | D2794 | 06-10-2017 | Kenton-on-Sea | 1;<br>ND;<br>*1;* | Potentially new species. Also present in France |
| Corallinaceae sp. 1 | - | D2826 | 07-10-2017 | Port Alfred | **1**;<br>ND;<br>ND; | Morphologically identified as *Jania verrucossa* but barcodes are not available for this gene. The closest relative is *Bossiella*, comparisons with other gene regions for *Bossiella* show a distant |

| | | | | | | |
|---|---|---|---|---|---|---|
| | | | | | | relationship between *Jania* and *Bossiella* which suggest this is probably a new species |
| Gelidiales<br>Gelidiaceae<br>*Gelidium abbottiorum* R.E.Norris | Widenham, IWP-MP, South Africa | D2771 | 06-10-2017 | Kenton-on-Sea | **1**;<br>*1;*<br>**1**; | New barcodes for an endemic species (DNA confirmation in EC) |
| *Gelidium pristoides* (Turner) Kützing | False Bay, BMP, South Africa | D2776;<br>D2858 | 06-10-2017;<br>09-10-2017 | Kenton-on-Sea;<br>Kenton-on-Sea | *2;*<br>2;<br>2; | |
| *Gelidium pteridifolium* R.E.Norris, Hommersand & Fredericq | Port Edward, IWP-MP, South Africa | D2821 | 07-10-2017 | Port Alfred | ND;<br>ND;<br>**1**; | New barcode for Southern African endemic |
| *Gelidium reptans* (Suhr) Kylin | Cape of Good Hope, BMP | D2916 | 29-05-2018 | Three Sisters | **1**;<br>ND;<br>1; | New barcodes based on specimens from South Africa |
| Gigartinales<br>Caulacanthaceae<br>*Caulacanthus* sp. 1 | - | D2820 | 07-10-2017 | Port Alfred | **1**;<br>1;<br>**1**; | SA material resolves within Lineage 2 according to Yang and Kim (2023) but is divergent from the type material |
| *Heringia mirabilis* (C.Agardh) J.Agardh* | Cape of Good Hope, BMP, South Africa | D2899 | 28-05-2018 | Port Alfred | **1**;<br>ND;<br>ND; | New barcode for Southern African endemic and monospecific species. Confirmed in the Gigartinales |
| Cystocloniaceae<br>*Calliblepharis fimbriata* (Greville) Kützing | Cape, BMP, South Africa | D2827 | 07-10-2017 | Port Alfred | **1**;<br>1;<br>ND; | New barcode based on SA material |
| Cystocloniaceae sp. 1 | - | D2795;<br>D2812;<br>D2927 | 06-10-2017;<br>07-10-2017;<br>30-05-2018 | Kenton-on-Sea;<br>Port Alfred;<br>Great Fish River | **2**;<br>**3**;<br>**3**; | Potentially new species and new genus |

| | | | | | | |
|---|---|---|---|---|---|---|
| *Hypnea* sp. 1 | Mtwalume, IWP-MP, South Africa | D2784 | 06-10-2017 | Kenton-on-Sea | **1**;<br>*1;*<br>ND; | Consistent with *Hypnea rosea* sp. 1 RSA Johnson 2017 |
| *Hypnea spicifera* (Suhr) Harvey | Algoa Bay, AMP, South Africa | D2772 | 06-10-2017 | Kenton-on-Sea | **1**;<br>**1**;<br>**1**; | New barcodes for the species from South Africa |
| *Tenebris* sp. 1 | - | D2800 | 06-10-2017 | Kenton-on-Sea | **1**;<br>**1**;<br>ND; | Consistent with *Tenebris* sp. (same as D1046), a new genus identified in Johnson 2017 |
| Gigartinaceae<br>*Chondracanthus acicularis* (Roth) Fredericq | Adriatic Sea | D2799 | 06-10-2017 | Kenton-on-Sea | 1;<br>ND;<br>1; | First barcodes for SA specimens confirms the presence of the species in the region |
| *Gigartina paxillata* Papenfuss | Storms River, AMP, South Africa | D2768 | 06-10-2017 | Kenton-on-Sea | **1**;<br>**1**;<br>**1**; | *G. polycarpa* and *G. paxillata* have non-overlapping distributions in SA. *G. polycrpa* is a typical west-coast species. The available sequence for the species collected from the east-coast is likely a misidentification as it resolved with *G. paxillata*. New barcodes for an SA endemic |
| *Gigartina pistillata* (S.G.Gmelin) Stackhouse | Doubtful | D2923; D2844 | 30-05-2018; 09-10-2017 | Great Fish River; Kenton-on-Sea | 2;<br>*2;*<br>1; | |
| Phyllophoraceae<br>*Gymnogongrus tetrasporifer* Papenfuss* | Needs designation of type locality | D2833 | 09-10-2017 | Kenton-on-Sea | **1**;<br>**1**;<br>ND; | Species described in an unpublished manuscript by Papenfuss but needs to have a designated type |
| *Gymnogongrus* sp. 1 | - | D2926 | 30-05-2018 | Great Fish River | **1**;<br>**1**;<br>ND; | Potentially new species |
| Rhizophyllidaceae<br>*Portieria hornemannii* (Lyngbye) P.C.Silva | Red Sea | D2897 | 28-05-2018 | Port Alfred | **1**;<br>1;<br>1; | |

| | | | | | | |
|---|---|---|---|---|---|---|
| Gracilariales<br>Gracilariaceae<br>Gracilarieae | | | | | | |
| *Gracilaria aculeata* (Hering) Papenfuss | Durban, IWP-MP, South Africa | D2781 | 06-10-2017 | Kenton-on-Sea | **1**;<br>**1**;<br>**1**; | New barcodes for a SA endemic. Resolves in the subgenus *Corallopsis* (Gurgel et al. 2018) |
| *Gracilaria beckeri* (J.Agardh) Papenfuss | Port Alfred, AMP, South Africa | D2770 | 06-10-2017 | Kenton-on-Sea | **1**;<br>1;<br>ND | New barcode based on SA specimen |
| *Gracilaria capensis* F.Schmitz ex Mazza | Port Alfred, AMP, South Africa | D2773;<br>D2971 | 06-10-2017;<br>11-09-2018 | Kenton-on-Sea;<br>Port Alfred | **2**;<br>*2;*<br>**1**; | New barcodes from the type locality of the species |
| *Gracilaria denticulata* F.Schmitz ex Mazza | Port Alfred, AMP, South Africa | D2909 | 29-05-2018 | Three Sisters | **1**;<br>1;<br>**1**; | A specimen of this species collected from SA from was mislabelled as *G. huangii* (DQ296121.1) see Lin and De Clerck 2006 for details. New barcodes based on SA material |
| Halymeniales<br>Halymeniaceae | | | | | | |
| *Codiophyllum natalense* J.E.Gray | Durban, IWP-MP, South Africa | D2894 | 28-05-2018 | Port Alfred | **1**;<br>*1*;<br>ND; | New barcode for SA endemic |
| *Pachymenia cornea* (Kützing) Chiang | Cape Agulhas, BMP, South Africa | D2819 | 07-10-2017 | Port Alfred | ND;<br>1;<br>**1**; | New barcode based on SA material |
| *Pachymenia ulvoidea* (F.Schmitz) L.K.Russell | Port Alfred, AMP, South Africa | D2816 | 07-10-2017 | Port Alfred | **1**;<br>ND;<br>**1**; | New barcodes fro SA endemic |
| *Prionitis filiformis* Kylin* | California, USA | D2783 | 06-10-2017 | Kenton-on-Sea | **1**;<br>*1*;<br>ND; | SA barcodes are different from USA material. SA material will need a new name |
| Hildenbrandiales<br>Hildenbrandiaceae | | | | | | |

| | | | | | | |
|---|---|---|---|---|---|---|
| *Hildenbrandia lecannellieri* Hariot | South America and Australia | D2808 | 06-10-2017 | Kenton-on-Sea | **1**;<br>ND;<br>ND; | First barcode for the species confirms its placement in the family Hildenbrandiaceae and order Hildenbrandiales |
| Nemaliales | | | | | | |
| Galaxauraceae | | | | | | |
| *Dichotomaria diesingiana* (Zanardini) Huisman, J.T.Harper & G.W.Saunders | Durban, IWP-MP, South Africa | D2814 | 07-10-2017 | Port Alfred | *1*;<br>1;<br>**1**; | New barcode based on SA material |
| *Dichotomaria hommersandii* S.-L.Liu & Showe M.Lin | Port Alfred, AMP, South Africa | D2835 | 09-10-2017 | Kenton-on-Sea | **1**;<br>1;<br>ND; | New barcodes for SA endemic |
| *Dichotomaria* sp. 1 | - | D2837 | 09-10-2017 | Kenton-on-Sea | **1**;<br>**1**;<br>**1**; | Morphologically identified as *D. tenenera* but does not group with this species. Potentially new species |
| Scinaiaceae | | | | | | |
| *Nothogenia erinacea* (Turner) P.G.Parkinson* | Cape of Hope, BMP, South Africa | D2815 | 07-10-2017 | Port Alfred | ND;<br>*1*;<br>ND; | New barcode for the species from the EC. Distantly related to the generitype (93% rbcL) |
| *Scinaia capensis* (Setchell) Huisman* | Port Alfred, AMP, South Africa | D2824 | 07-10-2017 | Port Alfred | **1**;<br>1;<br>ND; | First barcode from the type locality of the species (other barcodes from Tristan). Distantly related to the type species for the genus |
| Plocamiales | | | | | | |
| Plocamiaceae | | | | | | |
| *Plocamium beckeri* F.Schmitz ex Simons | Port Edward, IWP-MP, South Africa | D2793 | 06-10-2017 | Kenton-on-Sea | ND;<br>ND;<br>1; | First barcodes for the species from near its type locality. Misapplied name (JF271603), see Reddy et al. 2023 |
| *Plocamium robertiae* F.Schmitz ex Mazza | Port Alfred, AMP, South Africa | D2912;<br>D2977 | 29-05-2018;<br>11-09-2018 | Three Sisters<br>Port Alfred | **2**;<br>**1**;<br>ND; | Species reinstatement Reddy et al. 2023 |
| *Plocamium suhrii* Kützing | Cape of Good Hope, IWP-MP, South Africa | D2903;<br>D2910;<br>D2913 | 28-05-2018;<br>28-05-2018;<br>28-05-2018; | Port Alfred;<br>Three Sisters;<br>Three Sisters | 3;<br>2;<br>2; | New barcodes for the species confirm its presence in Spain. *Plocamium raphelisianum* reduced to a synonym of *P. suhrii* Reddy et al. 2023 |

| | | | | | | |
|---|---|---|---|---|---|---|
| Rhodymeniales | | | | | | |
| Rhodymeniaceae | | | | | | |
| *Halopeltis* sp. 1SA | - | D2911 | 29-05-2018 | Three Sisters | **1**;<br>1;<br>ND; | Barcode does not match *B. madagascariensis* |
| *Rhodymenia capensis* J.Agardh | Cape of Good Hope, BMP, South Africa | D2914 | 29-05-2018 | Three Sisters | **1**;<br>*1;*<br>**1**; | New barcodes based on SA material |
| *Rhodymenia natalensis* Kylin | Isipingo beach, IWP-MP, South Africa | D2915 | 29-05-2018 | Three Sisters | **1**;<br>**1**;<br>**1**; | First barcodes for a Southern African endemic species confirms its placement in *Rhodymenia* |

Key

ND=not determined

Bold=new barcode

Italics=new barcode for the EC

*=flagged for taxonomic revision
